# An adapted MS2-MCP system to visualize endogenous cytoplasmic mRNA with live imaging in *Caenorhabditis elegans*

**DOI:** 10.1101/2023.06.13.544769

**Authors:** Cristina Tocchini, Susan E. Mango

## Abstract

Live imaging of RNA molecules constitutes an invaluable means to track the dynamics of mRNAs, but live imaging in *Caenorhabditis elegans* has been difficult to achieve. Endogenous transcripts have been observed in nuclei, but endogenous mRNAs have not been detected in the cytoplasm, and functional mRNAs have not been generated. Here, we have adapted live imaging methods to visualize mRNA in embryonic epithelial cells. We have tagged endogenous transcripts with MS2 hairpins in the 3’ Untranslated Region (UTR) and visualized them after adjusting MS2 Coat Protein (MCP) expression. A reduced number of these transcripts accumulate in the cytoplasm, leading to loss-of-function phenotypes. In addition, mRNAs for *dlg-1* fail to associate with the adherens junction, as observed for the endogenous mRNA. These defects are reversed by inactivating the nonsense-mediated decay pathway. RNA accumulates in the cytoplasm, *dlg-1* associates with the adherens junction, and mutant phenotypes are rescued. These data suggest that MS2 repeats can induce the degradation of endogenous targets and alter the cytoplasmic distribution. Although our focus is RNAs expressed in epithelial cells during morphogenesis, this method can likely be applied to other cell types and stages.

**Summary statement:** An adapted MS2-MCP method to tag endogenous transcripts in *C. elegans* embryos for live imaging without affecting mRNA stability.

## Introduction

RNA molecules have a highly dynamic life. Through the different stages of their life, from synthesis to translation, and ultimately to degradation, RNAs move between different subcellular compartments and localize to specific subcellular organelles. Tracking an mRNA in real-time is pivotal to studying RNA dynamics, which cannot be achieved with fixed samples. Repeat copies of bacteriophage MS2 RNA hairpins have been used extensively to tag RNAs for live imaging (Bertrand et al., 1998; Jaramillo et al., 2008; Ng et al., 2011; Schmidt et al., 2020). The system allows the detection of single mRNA molecules in real-time and with high resolution, thanks to high affinity binding of fluorescently-labeled MS2-coat protein (MCP) to MS2 hairpins (Bertrand et al., 1998). This bacteriophage-derived, live-imaging approach has led to a general understanding of the dynamics during the different steps of the complex life of an mRNA, notably transcription, nuclear export, subcellular localization, and degradation (Bertrand et al., 1998; Heinrich et al., 2017; Lucas et al., 2013; Tutucci et al., 2018; Wells et al., 2007.). Recent studies have demonstrated the versatility of this system to tackle even more detailed steps of RNA metabolism (Halstead et al., 2016; Voigt et al., 2019). With the advent of the CRISPR/Cas9 technique, it has become possible to genetically engineer MS2 sequences at the endogenous locus. In this way, endogenous and not reporter transcripts are detected allowing a more precise description of what occurs in a cell at the physiological level.

In *C. elegans*, active sites of transcription have been visualized via a genetic trick that tracks fluorescently tagged NRDE-3 in nuclei (Toudji-Zouaz et al., 2021). The MS2-MCP system has also been used to visualize nascent RNAs derived from single-copy, integrated transgenes, and this approach has allowed visualization of transcriptional bursting in the germ line (Lee et al., 2019). Over-expressed mRNAs generated from multicopy extrachromosomal arrays have enabled the visualization of RNA dynamics in somatic cells (Li et al., 2021). Although promising, these examples did not generate functional RNAs from the endogenous locus, raising the question of whether the dynamics of transgene RNAs reflect endogenous regulation (Li et al., 2021). The lack of a system to visualize functional, endogenous transcripts live has been a bottleneck to advancing studies in RNA localization in *C. elegans*.

Here, we establish MS2-MCP tagging to visualize endogenous, functional transcripts for live imaging in *C. elegans*. The adaptation of the system for *C. elegans* consists of three modifications: i) a genetic background lacking a proficient nonsense-mediated decay (NMD) pathway to avoid degradation of MS2-tagged transcripts; ii) fluorescently-tagged MCP lacking a nuclear localization signal (NLS) to avoid aberrant sequestration of mRNAs in nuclei; iii) a weak promoter to maintain low levels of MCP. These adjustments allow tracking of endogenous transcripts and should prove useful for understanding the dynamics of RNA regulation.

## Results

### Insertion of a 24xMS2 sequence in the 3’UTR of an endogenous *C. elegans* gene may cause gene-specific developmental defects and aberrancies at the RNA level

The aim of our work was to adapt the MS2-MCP method to visualize functional, endogenous mRNAs with live imaging. It was not known why the MS2-MCP system had failed in the past to allow visualization of endogenous mRNA in the cytoplasm. To understand whether the presence of MS2 hairpins in endogenous 3’UTRs was the bottleneck for using MCP-MS2 in *C. elegans*, we inserted MS2 hairpins in two endogenous genes, *spc-1* and *dlg-1* (Fig. 1A-B). Using CRISPR/Cas9 (Ghanta et al., 2021; Ghanta & Mello, 2020), we inserted 24 MS2 hairpins (Bertrand et al., 1998; Tingey et al., 2022) in the endogenous 3’UTR sequence of available GFP-tagged alleles for the two genes. MS2 sequences were inserted separately in the first (“MS2 v1”, orange) or second half (“MS2 v2”, blue) of the endogenous (“endo”, black) 3’UTRs of *spc-1* (Fig. 1A) and *dlg-1* (Fig. 1B). PCR and sequencing analysis confirmed the location and sequence of our insertions.

**Figure 1.**
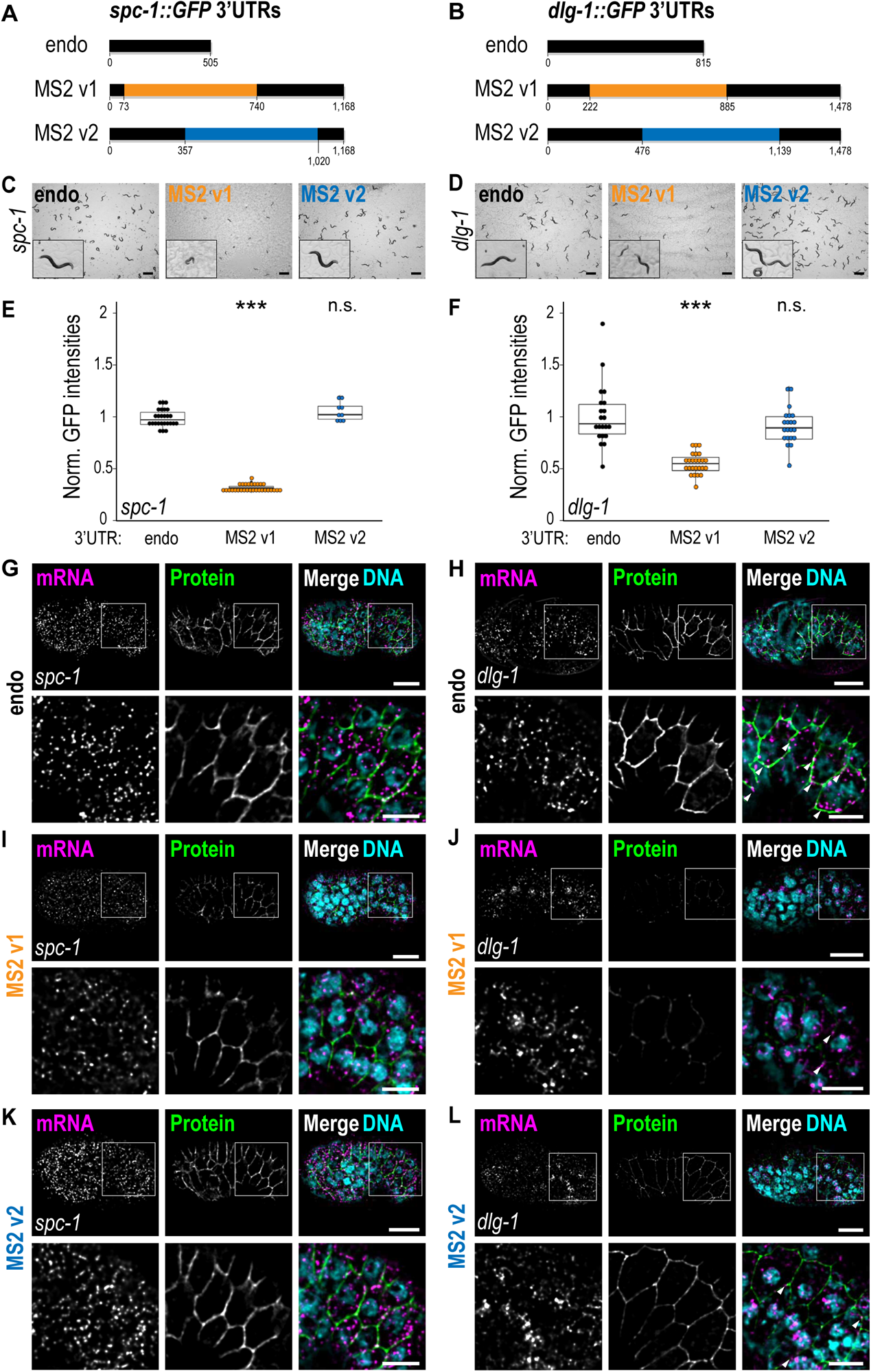
Insertion of MS2 hairpins in endogenous *C. elegans* 3’UTRs determines knock-down-like phenotypes. **A-B.** Schematic representations of the different versions of *spc-1* (**A**) and *dlg-1* (**B**) 3’UTRs used in this study. Untagged endogenous 3’UTRs are represented in black (“endo”) and their corresponding nucleotide length. The 24 MS2 hairpin insertions are either colored in orange for version 1 (“MS2 v1” – inserted in the first half of each 3’UTR), or in blue for version 2 (“MS2 v2” – inserted in the second half of each 3’UTR). The location of insertion and the final 3’UTR lengths are provided. **C-D.** Live images of free-living *C. elegans* animals on agar plates for the different 3’UTR strains of *spc-1* (**C**) and *dlg-1* (**D**). At the bottom-left corner of each image, a 10x magnified animal from each plate exemplifies the generally observed phenotypes. **E-F.** Dot plot with box plot: each dot represents the normalized GFP intensity of heads (*spc-1*) or pharynges (*dlg-1*) of young adult animals (24 hours after the L4 stage) with untagged endogenous (black), MS2 v1 (orange), and MS2 v2 3’UTRs (blue) of *spc-1* (**E**) and *dlg-1* (**F**). Significance of statistical analyses (*t*-test, two tails): n.s. > 0.05; *** < 0.001. **G-L.** Fluorescent micrographs of lateral views of *C. elegans* embryos at the comma stage (upper panels) carrying untagged endogenous (**G-H**), MS2 v1 (**I-J**), or MS2 v2 (**K-L**) 3’UTRs for the corresponding gene. The lower panels show zoom-ins from the portion of the seam cells highlighted in the upper panels with a white square. Panels from left to right: smFISH signal of mRNAs (magenta), fluorescent signal of GFP-tagged proteins (green), and merges with DNA (cyan). Arrowheads: examples of laterally localized *dlg-1* mRNAs. Scale bar: 10 μm (upper panels) and 5 μm (lower panels).

A survey of phenotypes revealed differences between MS2 insertion MS2 v1 vs MS2 v2 for both *spc-1* and *dlg-1*. The MS2 v1 insertions induced gene-specific developmental defects in homozygotes of both strains. For *spc-1,* MS2 v1 homozygous animals exhibited slow growth, lack of coordination (Unc), small body size (Sma), and low brood sizes (Fig. 1C). *dlg-1* MS2 v1 homozygous animals displayed a variable developmental delay resulting in an asynchronous population but were otherwise fertile and healthy (Fig. 1D). The observed phenotypes for both *spc-1* and *dlg-1* MS2 v1 strains resembled those deriving from partial inactivation of each gene (Kamath & Ahringer, 2003; Riga et al., 2021). Consistent with this idea, GFP imaging and quantitation revealed that *spc-1* and *dlg-1* MS2 v1 animals exhibited a significant decrease in protein (at least 50%) compared to animals carrying an endogenous 3’UTR (Fig. 1E-F, I-J).

In contrast to the MS2 v1 insertions, the MS2 v2 alleles had minimal phenotypic defects. *spc-1* MS2 v2 animals did not show any apparent developmental defects and were comparable to the untagged endogenous 3’UTR for both phenotype and protein expression levels (Fig. 1C, E, K). *dlg-1* MS2 v2 had a slight delay compared to the wild-type controls, but milder than *dlg-1* MS2 v1 (Fig. 1D). In line with this observation, protein levels in *dlg-1* MS2 v2 animals were decreased roughly 10% compared to the *dlg-1* endogenous 3’UTR strain (Fig. 1F, L).

We performed smFISH (Raj et al., 2008; Tsanov et al., 2016) to determine if the presence of the MS2 inserts caused problems at the RNA level. We found that all MS2 strains except *spc-1* MS2 v2 had a marked reduction in the number of cytoplasmic RNAs compared to their untagged controls (Fig. 1G-L). On the other hand, in nuclei, the *dlg-1* MS2 strains had greater signal intensities compared to the untagged endogenous 3’UTR strain (Fig. 1J, L) (see also next section). We also noted that the normal localization of *dlg-1* mRNA to adherens junctions was impaired in MS2 v1 (Fig. 1H) and, to a lesser extent, MS2 v2 (Fig. 1J, L). Overall, these data reveal that MS2 hairpins in the 3’UTR of *C. elegans* genes can interfere with normal mRNA accumulation and localization in the cytoplasm.

### An NMD-deficient background rescues the phenotypes determined by MS2 insertions in a gene 3’UTR

To begin to understand the defects induced by MS2 insertion, we examined the *spc-1* MS2 v2 strain, which did not exhibit any obvious defects. Endogenous *spc-1* carries a putative and previously unreported alternative polyadenylation (APA) signal at position 295 of the 3’ UTR, located upstream of the insertion site of MS2 v2 (Fig. 2A, green arrow). cDNA analyses by 3’ rapid amplification of cDNA ends (RACE) revealed that this APA signal was not used in the wild-type strain, but became the preferential one for *spc-1* MS2 v2, producing a 3’UTR in the mRNA that lacked the MS2 insert (Fig. 2A). We surmise that this truncated RNA produces wild-type levels SPC-1 protein and healthy animals, but would not be useful for live imaging.

**Figure 2.**
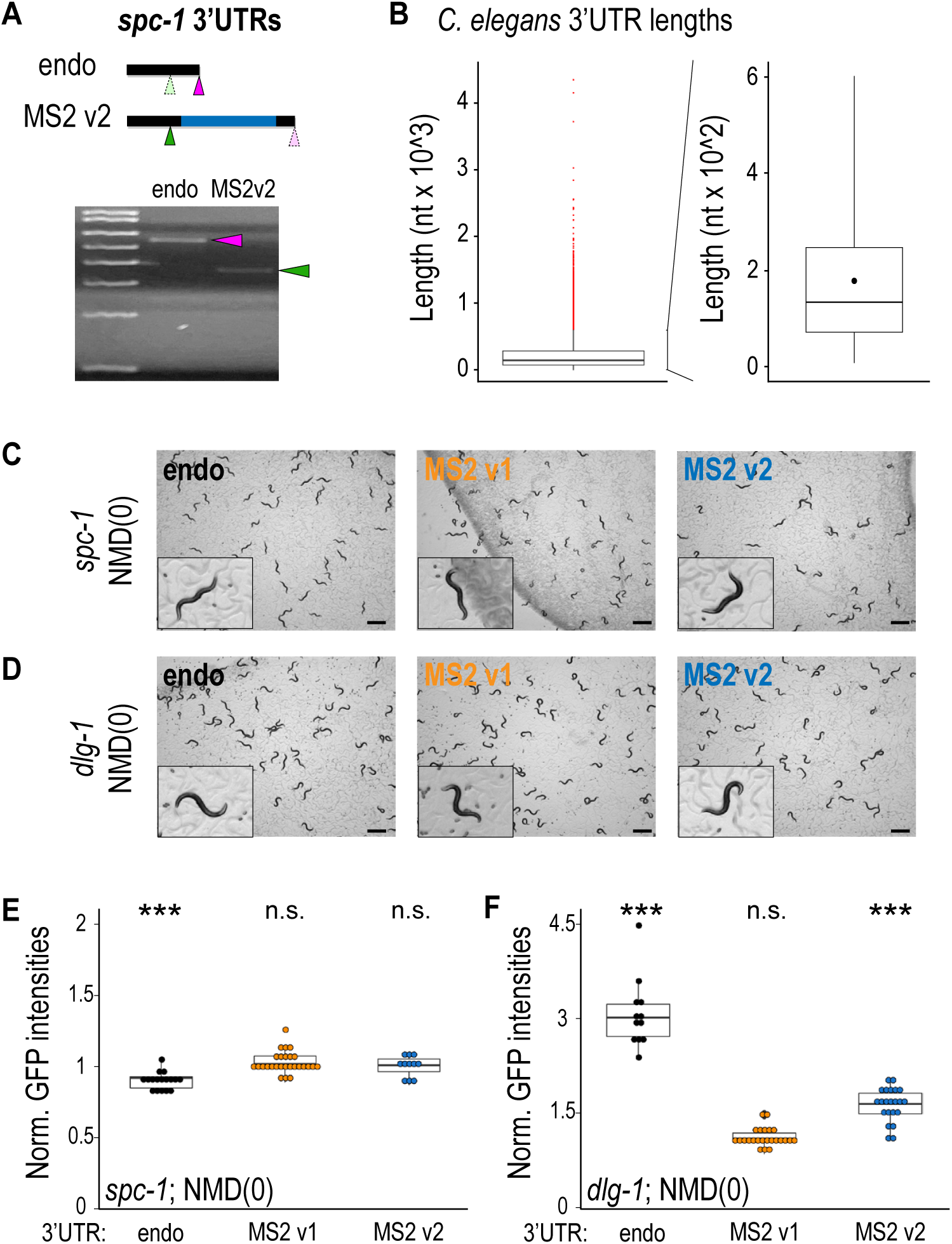
Alteration of the NMD pathway rescues the phenotypes caused by the MS2 insertion in endogenous *C. elegans* 3’UTRs. **A. B.** Left side: box plot of 3’UTR lengths (nucleotides) in *C. elegans*. Red dots: outliers. Right side: magnification of the box plot on the right, excluding the outliers. **C-D.** Live images of free-living *C. elegans* animals on agar plates for the different 3’UTR strains of *spc-1* (**C**) and *dlg-1* (**D**) in an NMD-deficient (NMD(0)) background. At the bottom-left corner of each image, a 10x magnified animal from each plate exemplifies the generally observed phenotypes. **E-F.** Dot plot with box plot: each dot represents the normalized GFP intensity of heads (*spc-1*) or pharynges (*dlg-1*) of young adult animals (24 hours after the L4 stage) with endogenous (black), MS2 v1 (orange), and MS2 v2 3’UTRs (blue) of *spc-1* (**E**) and *dlg-1* (**F**) in an NMD(0) background. Significance of statistical analyses (*t*-test, two tails): n.s. > 0.05; *** < 0.001.

Next, we examined the three remaining tagged strains, which each exhibited loss-of-function phenotypes. Prior studies have shown that mRNAs with artificial or endogenous but long 3’UTRs are destabilized by the NMD pathway in several systems (Bühler et al., 2006; Eberle et al., 2008; Hogg & Goff, 2010; Mango, 2001), suggesting that the transcripts with the MS2 hairpins might be targeted by NMD. The inserted MS2 sequence was 663-base pairs long, which roughly doubled the length of the *spc-1* and *dlg-1* 3’UTRs. *C. elegans* 3’UTRs are relatively short compared to other organisms, and they possess a mean of 223 nucleotides and a maximum of 598 nucleotides, excluding a few outliers (Fig. 2B; (Davis et al., 2022)). With the MS2 sequence, *spc-1* and *dlg-1* 3’UTRs reached a final length of 1,168 and 1,478 nucleotides, respectively (Fig. 1A-B), making them abnormally long.

To test if the presence of the MS2 sequence could destabilize transcripts via NMD (Fig. 1G-L), we introduced the strains for both *spc-1* and *dlg-1* carrying either the untagged endogenous gene or the MS2 tags into an NMD-deficient background (“NMD(0)”). All the morphological and developmental abnormalities previously observed in the MS2-tagged strains for both *spc-1* and *dlg-1* (Fig. 1C-D) were rescued in the absence of a proficient NMD pathway, and the animals were comparable to their respective controls (Fig. 2C-D). In line with the phenotypic rescue, analysis of protein abundance revealed that the GFP levels were increased for all the strains in the NMD(0) background. For *dlg-1* MS2 v2, levels were equivalent to the untagged 3’UTR in the wild-type background (Fig. 2E-F). The protein levels for *dlg-1* untagged and for MS2 v2 in the NMD(0) background were significantly increased compared to endogenous *dlg-1* in the wild-type background (Fig. 2F). *dlg-1* transcript contains a remarkably long 3’UTR (815 nucleotides) for *C. elegans* mRNAs (Fig. 2B). Although transcripts possessing long 3’UTRs can evade NMD-mediated degradation in some instances (Eberle et al., 2008; Toma et al., 2015), alteration of the NMD pathway would not affect the levels of the protein product of such transcripts. This is in contrast to what we observed for the endogenous *dlg-1* mRNA (Fig. 2F). We conclude that MS2 hairpins render *dlg-1* and *spc-1* targets of NMD and that, in addition, *dlg-1* is a *bona fide* endogenous target of the NMD pathway that has not been previously reported (Muir et al., 2018).

We wanted to verify that a deficiency in the NMD pathway could rescue the observed phenotypes by preventing RNA degradation and the other consequent RNA defects observed in the MS2 strains (*e.g.*, reduced cytoplasmic RNA levels, loss of subcellular transcript localization, and nuclear RNA clusters). To test this idea, we performed smFISH experiments on the different *dlg-1* strains in the presence or absence of a functional NMD pathway.

smFISH for the untagged, endogenous *dlg-1* 3’UTR strains in wild-type and NMD(0) backgrounds showed that impairment of the NMD pathway did not cause any defects at the level of mRNA abundance or localization for *dlg-1* transcripts (Fig. 3A) (Tocchini et al., 2021). We conclude that the NMD pathway is not critical for endogenous *dlg-1* regulation. However, the NMD-deficient background had profound effects on the tagged RNAs. It was sufficient to rescue both cytoplasmic RNA abundance and subcellular transcript localization at adherens junctions for both *dlg-1* MS2 v1 and MS2 v2 strains (Fig. 3B-C). These data reveal that active NMD can interfere with MS2-tagged RNA localization (see Discussion).

**Figure 3.**
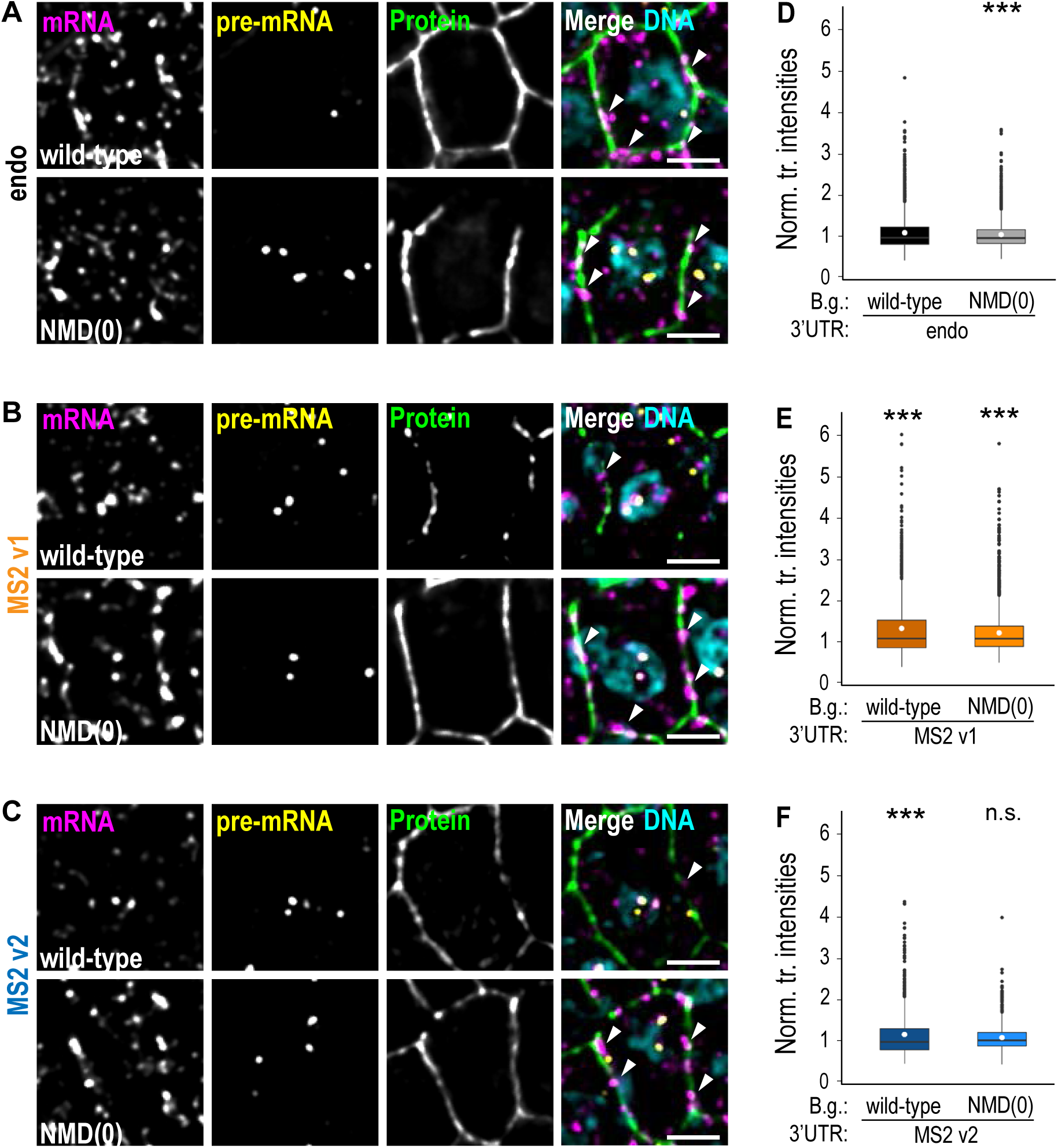
Alteration of the NMD pathway rescues the RNA phenotypes caused by the MS2 insertion in 3’UTRs *dlg-1* 3’UTR. **A-C.** Fluorescent micrographs of examples of seam cells of *C. elegans* embryos at the bean stage carrying a native (black, (**A**)), an MS2 v1 (orange, (**B**)), or an MS2 v2 (blue, (**C**)) 3’UTR in a wild-type (upper panels) or in a NMD(0) (lower panels) background. Panels from left to right: smFISH signal of endogenous GFP-tagged *dlg-1* mRNAs (magenta), smFISH signal of endogenous GFP-tagged *dlg-1* pre-mRNAs (yellow), fluorescent signal of GFP-tagged DLG-1 (green), and merges with DNA (cyan). Arrowheads: examples of laterally localized mRNAs. Scale bar: 2.5 μm. **D-F.** Dot plot with box plots: each dot represents the normalized smFISH fluorescent intensity detected with *dlg-1* intron probes from nuclei of whole embryos at the bean stage (n = 5 for each strain). The data derive from embryos with a native 3’UTR in the wild-type (“wild-type”; mean = 1.00; median = 0.87; StDev = 0.46) or in the NMD-deficient background (“B.g.”) (“NMD(0)”; mean = 0.95; median = 0.86; StDev = 0.37), in black (**D**); an MS2 v1 3’UTR in the wild-type (mean = 1.27; median = 1.00; StDev = 0.79), or in the NMD(0) background (mean = 1.15; median = 1.00; StDev = 0.56), in orange (**E**); an MS2 v2 3’UTR in the wild-type (mean = 1.10; S.E.M. = 0.02), or in the NMD(0) background (mean = 1.01; median = 0.94; StDev = 0.35), in blue (**F**). Significance of statistical analyses (*t*-test, two tails): n.s. > 0.05; *** < 0.001.

The smFISH experiments performed with exonic smFISH probes showed bright RNA dots within nuclei in the *dlg-1* MS2 strains compared to the untagged endogenous 3’UTR control (Fig. 1H, J, L and Fig. 3A-C). The brightness of smFISH dots correlates with the number of transcripts located in the immediate vicinity (Lee et al., 2019). Such nuclear RNA clusters could therefore represent the upregulation of transcription to compensate for the deleterious effects of the MS2 inserts, somewhat reminiscent of what has been described as transcriptional adaptation (Serobyan et al., 2020). smFISH experiments with intronic smFISH probes revealed colocalization of the detected *dlg-1* pre-mRNA with the previously described nuclear clusters, demonstrating that they represented transcriptional sites (Fig. 3A-C). Quantitation of the fluorescence intensities of the transcriptional sites stained with smFISH intron probes revealed an increase in the transcriptional output in both *dlg-1* MS2 strains in the wild-type background compared to the control (Fig. 3D-F). When lacking an efficient NMD pathway, the transcriptional levels were partially (MS2 v1) or fully (MS2 v2) restored to the levels of the untagged, endogenous 3’UTR strain in the wild-type background (Fig. 3D-F). These results demonstrated the existence of an inverse correlation between DLG-1 protein levels and *dlg-1* transcriptional levels, possibly reflecting the existence of a feedback mechanism to control DLG-1 protein levels through transcription. In summary, the lack of a proficient NMD pathway largely restored the defects for MS2 v1 and v2, including developmental phenotypes, reduced protein and transcript levels, loss of subcellular mRNA localization, and increased transcriptional levels.

### Lowly expressed fluorescently-labeled MCP lacking an NLS allows the visualization of mRNA in live *C. elegans* embryos

Previous studies used MCP tagged with an NLS to reduce background signal in the cytoplasm (Bertrand et al., 1998). We reasoned that the use of an NLS-tagged MCP might interfere with the proper processing or nuclear export of an endogenous mRNA. To circumvent such deleterious effects, we designed a cytoplasmic MCP that lacked an NLS. Specifically, we generated a single-copy, integrated transgene where two MCPs were tagged with FLAG and mCherry sequences (designated “MCP”) and expressed under the control of a heat-shock promoter (Fig. 3A). Heat-shock promoters possess a low activity at physiological temperatures of 25°C in somatic cells (Tocchini et al., 2014) which allows the production of low levels of the desired protein. The low temperature reduces the surplus of cytoplasmic MCP that was previously achieved by sequestering unbound MCP in nuclei.

To determine if the combination of MS2-tagged RNA, with cytoplasmic MCP, and the NMD(0) background could recapitulate the behavior of endogenous transcripts in wild-type animals, we performed live imaging on the strains carrying MS2 v1 3’UTRs. Both *spc-1* and *dlg-1* MS2 v1 cytoplasmic transcripts could be detected with fluorescently-tagged MCP in the NMD(0) background at the same stages and in the same cell types as their untagged endogenous counterparts (Fig. 4B-C). Importantly, using this configuration, *dlg-1* transcripts were localized at adherens junctions (Fig. 4C, white arrowheads). Although heat-shock promoters are ubiquitously expressed in somatic embryonic cells, we noticed different extents of MCP expression between and within embryos grown at 25°C (Fig. 4C, white asterisks). Some embryos did not express MCP at all, whereas others expressed it but not in all cells. Overall, our optimized MS2-MCP system paired with an NMD-deficient background allowed visualization of test transcripts in live *C. elegans* embryos.

**Figure 4.**
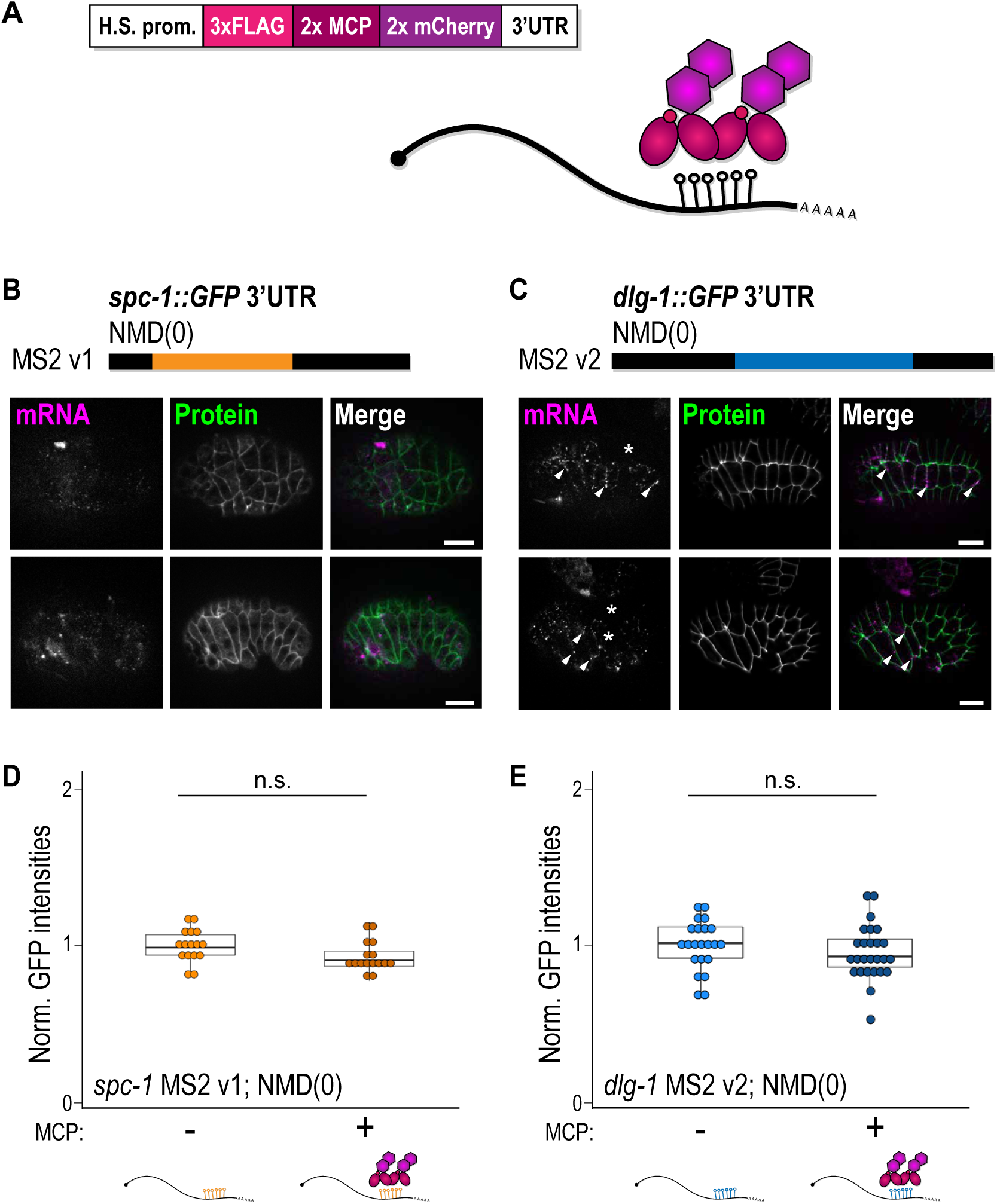
MS2-tagged transcripts can be visualized with live imaging thanks to a lowly expressed fluorescently-tagged MCP. **A.** Schematic representation of the MCP construct used in this study and of the MS2-MCP system. Two copies of MCP sequences (2x MCP) are fused to a 3xFLAG-tag sequence (3xFLAG) and two copies of mCherry sequences (2x mCherry). The expression of the transgene is under the control of the heat-shock promoter of the *hsp-16.48* gene (H.S. prom.), and the 3’UTR derives from the *tbb-2* gene (3’UTR). **B-C.** Schematic representations of the MS2 v1 of *spc-1* and v2 of *dlg-1* 3’UTRs in the NMD-deficient background used for live imaging. Live fluorescent images of *C. elegans* embryos: 8E (upper panels) and comma stages (lower panels) for *spc-1* (**B**), and bean (upper panels) and 1.5-fold stages (lower panels) for *dlg-1* (**C**). Panels from left to right: live signal of MS2 v1 mRNAs for *spc-1* (**B**) and *dlg-1* (**C**) visualized through the fluorescently labeled MCP (magenta), fluorescent signal of GFP-tagged proteins (green), and merges. Asterisks: examples of embryonic cells not expressing MCP. Arrowheads (**C**): examples of laterally localized *dlg-1* mRNAs. Scale bar: 10 μm. **D-E** Dot plot with box plot: each dot represents the normalized GFP intensity of heads (*spc-1*) (**D**) or pharynges (*dlg-1*) (**E**) of young adult animals (24 hours after the L4 stage) with MS2 v1 (*spc-1*) or v2 (*dlg-1*) 3’UTRs in the NMD-deficient background in the absence (“-”) or presence (“+”) of MCP (schematically represented underneath). Raw data in (**E**), minus are the same as in Fig. 2F as derived from the same experiments. Significance of statistical analyses (*t*-test, two tails): n.s. > 0.05.

The ectopic binding of a protein to a transcript’s 3’UTR can disrupt the translational regulation of the given mRNA (Szostak & Gebauer, 2013). Yet, no phenotypes were observed in NMD(0) animals engineered with the MS2-MCP system. To confirm that binding of MCP to the MS2-tagged mRNA did not affect translational output, we tested protein levels in the presence or absence of MCP in MS2-tagged strains in the NMD(0) background and also in the wild-type background for *dlg-1* MS2 strains. Analyses of protein levels through GFP quantification did not reveal a significant difference for the strains in the NMD(0) background in absence or presence of MCP (Fig. 4D-E; Fig. S1). In a wild-type background with a proficient NMD pathway, *dlg-1* MS2 v2 did also not show a significant difference in the two conditions tested with and without MCP, but *dlg-1* MS2 v1 showed a slight although significant increase in protein output in the presence of MCP compared to its absence (see discussion). Together, these results imply that MCP does not significantly affect the translational output of the transcript it binds at least in NMD(0). In conclusion, a lowly expressed and fluorescently-tagged MCP lacking an NLS can effectively bind transcripts tagged with MS2 hairpins and allow their visualization through live imaging, recapitulating endogenous mRNA and protein scenarios.

## Discussion

### MS2-tagged 3’UTRs are targeted by the NMD pathway

This study establishes the MS2-MCP system for live imaging of endogenous cytoplasmic transcripts in *C. elegans*. We showed that the presence of MS2 sequences in the endogenous 3’UTR of a *C. elegans* transcript can cause gene-specific loss of function phenotypes. Such phenotypes are mediated by the NMD pathway, suggesting that MS2-tagged mRNAs are destabilized by this surveillance mechanism. Inactivation of NMD combined with low level expression of fluorescently-labeled, cytoplasmic MCP enabled live imaging of cytoplasmic mRNA.

Our findings suggest that abnormally long 3’UTRs after MS2 insertion engenders post-transcriptional defects for the tagged mRNA. *C. elegans* 3’UTRs are relatively short compared to those of other systems, with a median length one-sixth of human 3’UTRs and a length distribution comparable to yeast (Jan et al., 2011). The length of the MS2 sequence alone goes beyond the upper limit of *C. elegans* 3’UTR length (Fig. 2B). Transcripts containing long 3’UTRs are recognized as aberrant by the NMD pathway (Bühler et al., 2006; Hogg & Goff, 2010; Mango, 2001), and the use of an NMD-deficient mutant can prevent their degradation. The physical distance between the polyadenylation (poly(A)) and the termination codon (TC) is responsible for whether a transcript becomes an NMD target (Eberle et al., 2008). Transcripts possessing long 3’UTRs have higher chances of finding their poly(A) and TC distant from each other. Nevertheless, such transcripts can still evade, at least to some extent, NMD-mediated degradation if their poly(A) and termination codon are in close proximity in the three-dimensional space (Eberle et al., 2008) (Fig. 5). We suggest that the increased length of the 3’UTR after insertion of MS2 hairpins likely explains why previous attempts to generate functional, tagged mRNAs failed.

**Figure 5.**
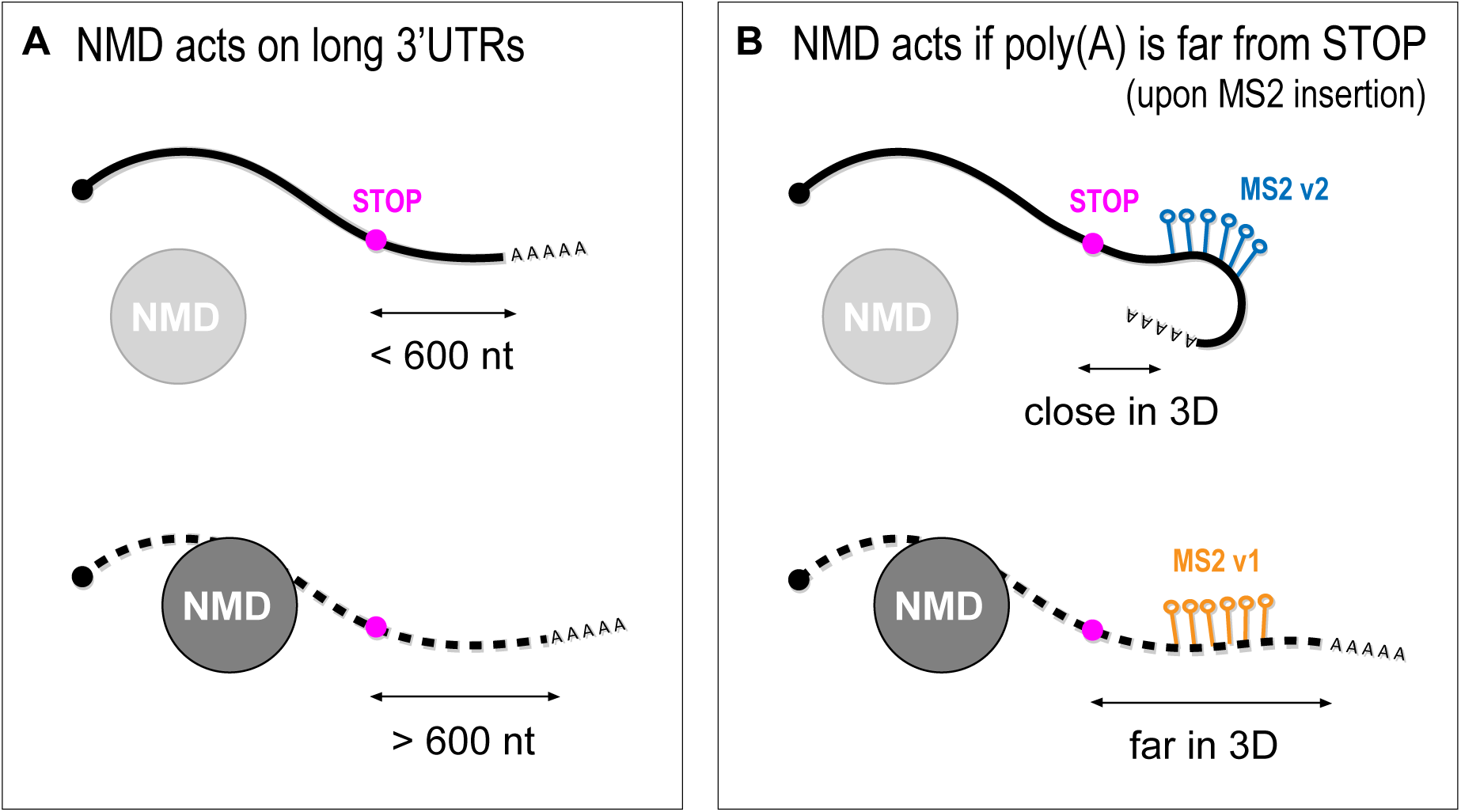
Long 3’UTRs are targeted by the NMD pathway. **A.** Schematic representation of transcripts with short (< 600 nucleotides) or long (> 600 nucleotides) 3’UTRs. NMD does not mediate the degradation of mRNA carrying a short 3’UTR but does degrade transcripts with long 3’UTRs. **B.** Schematic representation of the same transcript possessing MS2 hairpins inserted in two different sites of its 3’UTR. Both insertions determine the creation of an artificial long 3’UTR. In the v2 instance (blue hairpins), the poly(A) gets close to the STOP codon in the three-dimensional (3D) space, which prevents NMD to degrade the transcript. In the v1 instance (orange hairpins), the poly(A) is far from the STOP codon in the 3D space, and the mRNA is targeted by the NMD for degradation.

Our data show that the position of the MS2 hairpins within the 3’UTR matters. We observed stronger developmental phenotypes for *dlg-1* MS2 v1 compared to MS2 v2, although MS2 v1 and v2 transcripts possess the same MS2 sequence inserted in their 3’UTR. This oddity suggests that the NMD pathway can recognize MS2 v1 better or faster compared to v2. Based on the poly(A)-TC proximity model, we speculate that the different sites of insertion might cause distinct three-dimensional structures of the mRNA that allow (in MS2 v1) or not (in MS2 v2) the NMD pathway to act more or less strongly on them (Fig. 5). Alternatively, the distinction between v1 and v2 may reflect differential kinetics for degradation. For example, scanning from the stop codon or the poly(A) site might be differentially affected by MS2 hairpins in position v1 or v2. We note that in either model, MCP is not involved as its presence had little effect on the phenotypes at the organismal or protein levels. Only the position within the RNA was critical.

*C. elegans* animals that do not possess a functional NMD pathway can progress through all stages of development with only minor developmental defects (Mango, 2001; Pulak & Anderson, 1993). The NMD pathway plays a more critical role in higher eukaryotes, where its full removal induces major developmental defects or even lethality due to cytotoxicity (Usuki et al., 2013; Vicente-Crespo & Palacios, 2010). Nevertheless, we envision that a partial impairment of the NMD pathway may allow the visualization of MS2-tagged transcripts and therefore the method can be potentially extended to other systems, too. This partial inhibition can be achieved by gene silencing (Usuki et al., 2013) or inhibitory drugs (Zhao et al., 2022) that target key components of the NMD pathway.

### MCP: low levels and lack of an NLS

MS2-tagged transgenic transcripts can be visualized live with an MCP protein coupled to fluorophores (Bertrand et al., 1998). We removed the NLS that is usually included in MCP constructs and were able to visualize MS2-tagged RNAs using a weak promoter to reduce MCP levels. Low MCP levels were sufficient to visualize transcripts presumably due to the high affinity of MCP to MS2 hairpins, which implies a slow off-rate (Johansson et al., 1997), and guaranteed an optimal signal-to-noise ratio. Nevertheless, we observed some variability in MCP expression among and within embryos. If this represents a limitation, we recommend i) the use of a different but still lowly expressed promoter for MCP that can even be specific for the cells or tissue of interest, or ii) a mild heat-shock followed by a long recovery time to increase MCP levels still avoiding aberrant mRNA distribution (Tocchini et al., 2021).

In contrast to MS2 insertion *per se*, the binding of MCP to an MS2-tagged endogenous had little effect on the translational output (Fig. 4D-E; Fig. S1). Although the differences in protein levels were not significant in the absence or presence of MCP (besides *dlg-1* MS2 v1 in a wild-type background (Fig. S1)), it was possible to observe two general, although minor, trends: in a wild-type background, the presence of MCP increased protein output, whereas in an NMD(0) background, MCP lead to a decrease. We speculate that binding of MCP to the MS2-tagged mRNA might protect the mRNA from NMD-mediated degradation in a wild-type background. On the other hand, when the NMD pathway is inhibited, MCP might have a marginal role in post-transcriptional gene regulation via stabilizing the mRNA resulting in a higher protein output per transcript and overall.

In summary, we have developed an adapted MS2-MCP system for live imaging for the *C. elegans* model organism. This system overcame a previous limitation in the field, and it allows the visualization of endogenous transcripts tagged with MS2 in the cytoplasm through fluorescently-tagged MCP. The MS2 hairpins can be easily inserted in the genome via CRISPR thanks to their optimized and compact sequence (around 650 base-pairs). Differently from the recently reported MS2-based signal Amplification with Suntag System (MASS) (Hu et al., 2023), our method provides the possibility to pair the MS2-MCP system to the SunTag system to visualize translation to the same mRNA. We showed the system is able to recapitulate the subcellular localization of previously analyzed endogenous transcripts (*i.e.*, *dlg-1* at adherens junctions). This makes the system amenable for further studies to address how subcellular localization occurs and which pathway controls this phenomenon. We focused our study on developing embryonic epithelial cells, but our method has the potential to be extended to other cell types and developmental stages.

## Materials and methods

### Nematode culture

All animal strains were maintained as previously described (Brenner, 1974) at 20°C. Strains subjected to MS2-MCP live imaging were shifted to 25°C as young adults, and their progeny was imaged at embryonic stages specified in the figure legends a day after. For a full list of alleles and transgenic lines, see Table S1.

### Generation of transgenic lines

The 24xMS2 hairpin sequence was amplified from the plasmid pIE5 (gift from Jeffry Chao). CRISPR insertions were performed as previously described (Ghanta et al., 2021; Ghanta & Mello, 2020). Correct insertions were verified by genotyping and sequencing. For a full list of crRNAs and oligos used to amplify and validate the correct insertion of the 24xMS2 sequence, see Table S2.

The MCP transgene was built as follows (Fig. 4A): a *hsp-16.48* promoter (amplified from N2 lysate), a 3xFLAG tag (dsDNA oligo, ordered from IDT), 2xMCP sequences (amplified from the plasmid pIK270 - gift from Iskra Katic) paired to two mCherry sequences by an elastic linker (amplified from pCT2 and gBlock, ordered from IDT), and a *tbb-2* 3’UTR (amplified from N2 lysate) were assembled in the pCFJ150 vector to create the pCT3.37 plasmid (NEBuilder® HiFi DNA Assembly Cloning Kit, New England BioLabs, cat#E5520). The protocol previously described for the generation of mosSCI transgenic lines (Frøkjær-Jensen et al., 2008) was enrolled to integrate the MCP transgene on chromosome II (Table S1).

### RNA extraction and 3’RACE

Total RNA was extracted using the standard Phenol/Chloroform protocol from one nearly starved 6 cm plate with mixed-stage animals and eggs for each genotype. 3’RACE experiments were performed with the 5’/3’ RACE Kit, 2^nd^ Generation (Roche, cat#03353621001) for cDNA production paired to Phusion High-Fidelity DNA Polymerase (New England BioLabs, cat#M0530) for amplification. The oligo oCT3.471 (tcgccaattgtcatatgattt) was used as a forward primer to amplify the different 3’UTR segments.

### smFISH and live imaging

smFISH experiments were performed in duplicates, following the protocol described in (Tocchini et al., 2021). smFISH probes were designed as previously described (Tsanov et al., 2016) and ordered from IDT (Tocchini et al., 2021). For a full list of smFISH probes, see Table S3.

Live imaging experiments were performed on animals at the indicated stage. Pharyngeal GFP signals were quantified on 1-day-old animals as previously described (Tocchini et al., 2021) For MS2-MCP live imaging experiments, embryos were washed off from plates with 1 ml of water, collected into Eppendorf tubes, and spun-down (“short”) for 5 seconds. After removing all the water, M9 buffer was added and embryos were resuspended and transferred onto poly-lysine-coated slides (Thermo Scientific, cat#ER-308B-CE24) and sealed with a coverslip.

### Microscopy, image analysis, and quantitation

A widefield ZEISS Axio Zoom V16 equipped with a ZEISS Axiocam 503 mono camera, and a ZEN 2.6 software (blue edition) were used for capturing images of free-living animals on agar plates.

As previously described (Tocchini et al., 2021), a widefield microscope FEI “MORE” with total internal reflection fluorescence (TIRF), equipped with a Hamamatsu ORCA flash 4.0 cooled sCMOS camera, and a Live Acquisition 2.5 software were used for capturing smFISH images.

A widefield ZEISS Axio Imager M2 equipped with an ApoTome.2, a ZEISS Colibri as an LED light source, a Hamamatsu ORCA flash 4.0 camera, and a ZEN 2.6 software (blue edition) were used for capturing images for live imaging experiments.

smFISH pictures were deconvolved with the Huygens software. All images were processed in OMERO (https://www.openmicroscopy.org/omero/), and figures were assembled in Adobe Illustrator (https://www.adobe.com/).

TrackMate from the python software (https://www.python.org/) has been used to quantify the fluorescent intensity of transcriptional dots. Statistical analyses were performed with the R software (R Core Team, 2021; https://www.R-project.org/). The ggplot2 package (Wickham, 2016; https://ggplot2.tidyverse.org) in R was used to generate dot plots with box plots: a thick horizontal line represents the median, hinges for the first and third quartiles, and whiskers mark upper and lower limits. Statistical differences were defined by *t*-test. For raw data, see Table S1.

## Supporting information

Table S1_Raw data

## Acknowledgments

We thank Dr. Iskra Katic and Dr. Jeffry Chao from the Friedrich Miescher Institute for Biomedical Research for sharing their plasmids for MCP (pIK270) and MS2 (pIE5), the Imaging Core Facility of the Biozentrum for technical support, current and previous lab members of the Mango group for scientific discussions, and WormBase. A special thanks to Dr. Sébastien Herbert from the IMCF for developing the script to analyze transcriptional levels on smFISH images and Dr. Fei Xu from the Mango group for technical assistance. Some strains were provided by the CGC, which is funded by the NIH Office of Research Infrastructure Programs (P40 OD010440).

## Table legends

**Table 1.** Strain list. List of names and respective genotypes of the C. elegans strains and transgenic lines used in this study.

| Strain name | Genotype | Reference |
| --- | --- | --- |
| N2 | wild-type | Brenner, 1974 (CGC) |
| SM1910 | *smg-2(e2008)* | Domeier *et al.*, 2000 |
| EG6699 | *ttTi5605 II; unc-119(ed3) III; oxEx1578* | Frøkjær-Jensen *et al.*, 2008 |
| SM2734 | *pxSi24[hsp-16.48p::3xFLAG::2xMCP::2xmCherry::tbb-2 3'UTR, unc-119(+)] II; unc-119(ed9) III* | This study |
| GOU2936 | *spc-1(cas815[spc-1::GFP]) X* | Jia *et al.*, 2019 |
| SM2815 | *spc-1(px126[spc-1::GFP::MS2_v1]) X* | This study |
| SM2817 | *spc-1(px128[spc-1::GFP::MS2_v2]) X* | This study |
| ML2615 | *dlg-1(mc103[dlg-1::GFP]) X* | CGC |
| SM4496 | *dlg-1(px139[dlg-1::GFP::MS2_v1]) X* | This study |
| SM2801 | *dlg-1(px118[dlg-1::GFP::MS2_v2]) X* | This study |
| SM2856 | *smg-2(e2008) I; spc-1(cas815[spc-1::GFP]) X* | This study |
| SM2852 | *smg-2(e2008) I; spc-1(px126[spc-1::GFP::MS2_v1]) X* | This study |
| SM2841 | *smg-2(e2008) I; spc-1(px128[spc-1::GFP::MS2_v2]) X* | This study |
| SM2669 | *smg-2(e2008) I; dlg-1(mc103[dlg-1::GFP]) X* | This study |
| SM4497 | *smg-2(e2008) I; dlg-1(px139[dlg-1::GFP::MS2_v1]) X* | This study |
| SM2807 | *smg-2(e2008) I; dlg-1(px118[dlg-1::GFP::MS2_v2]) X* | This study |
| SM2857 | *smg-2(e2008) I; pxSi24[hsp::MCP::mCherry, unc-119(+)] II; spc-1(px126[spc-1::GFP::MS2_v1]) X* | This study |
| SM4527 | *pxSi24[hsp::MCP::mCherry, unc-119(+)] II; dlg-1(px139[dlg-1::GFP::MS2_v1]) X* | This study |
| SM2803 | *pxSi24[hsp::MCP::mCherry, unc-119(+)] II; dlg-1(px118[dlg-1::GFP::MS2_v2]) X* | This study |
| SM4498 | *smg-2(e2008) I; pxSi24[hsp::MCP::mCherry, unc-119(+)] II; dlg-1(px139[dlg-1::GFP::MS2_v1]) X* | This study |
| SM2805 | *smg-2(e2008) I; pxSi24[hsp::MCP::mCherry, unc-119(+)] II; dlg-1(px118[dlg-1::GFP::MS2_v2]) X* | This study |

**Table 2.**
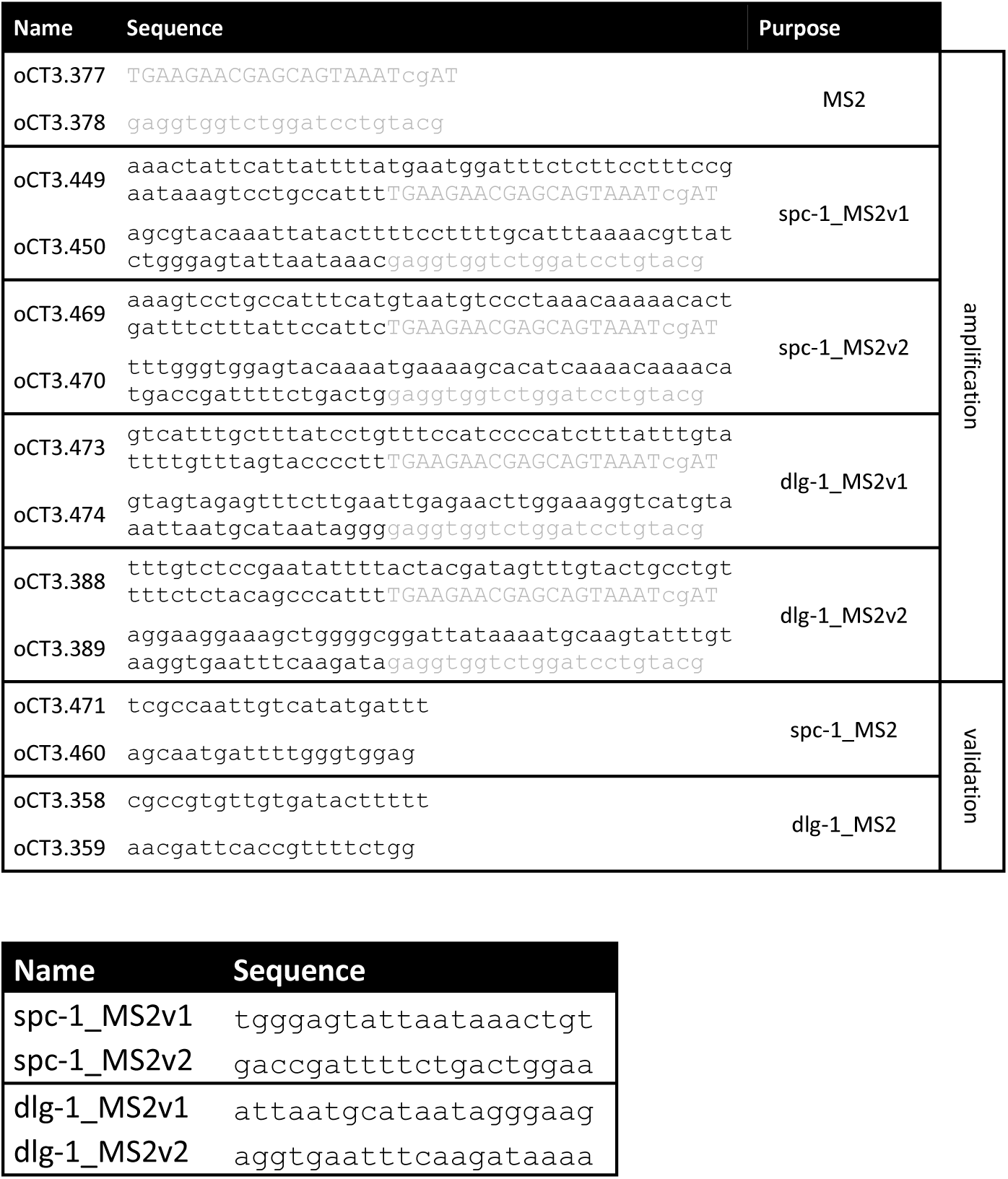
Oligos and crRNAs. List of oligos and crRNAs used in this study to generate and verify MS2 sequence insertion in CRISPR lines.

**Table 3.**
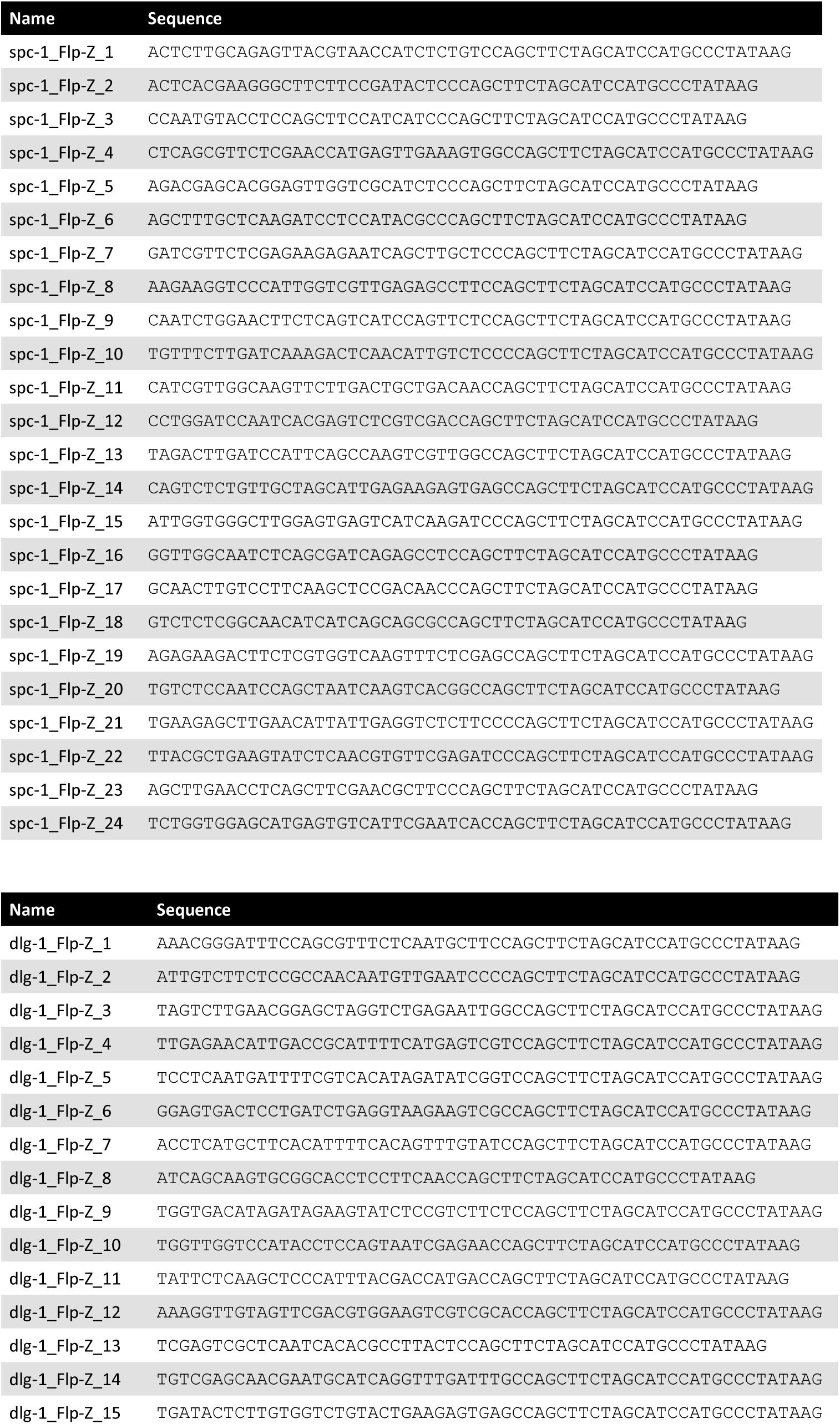
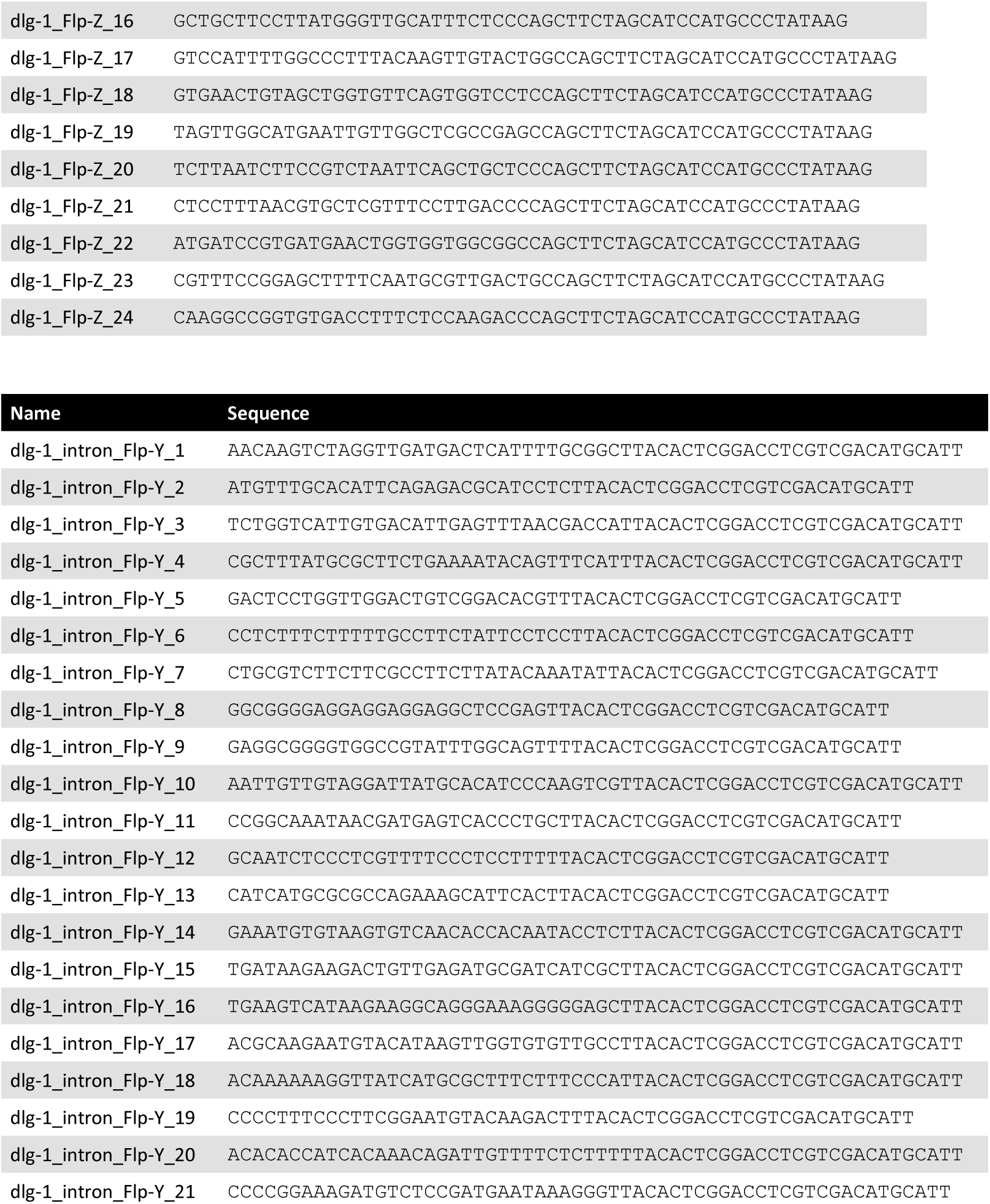
smFISH probes. List of smFISH probes used to detect spc-1 and dlg-1 mature RNAs, and dlg-1 nascent transcript (intron probes).

## Supplemental table legends

**Table S1. Raw data.** List of quantitation of GFP intensities of heads (*spc-1*) or pharynges (*dlg-1*) in the different figures.

## Supplemental figure legends

**Supplemental figure 1.**
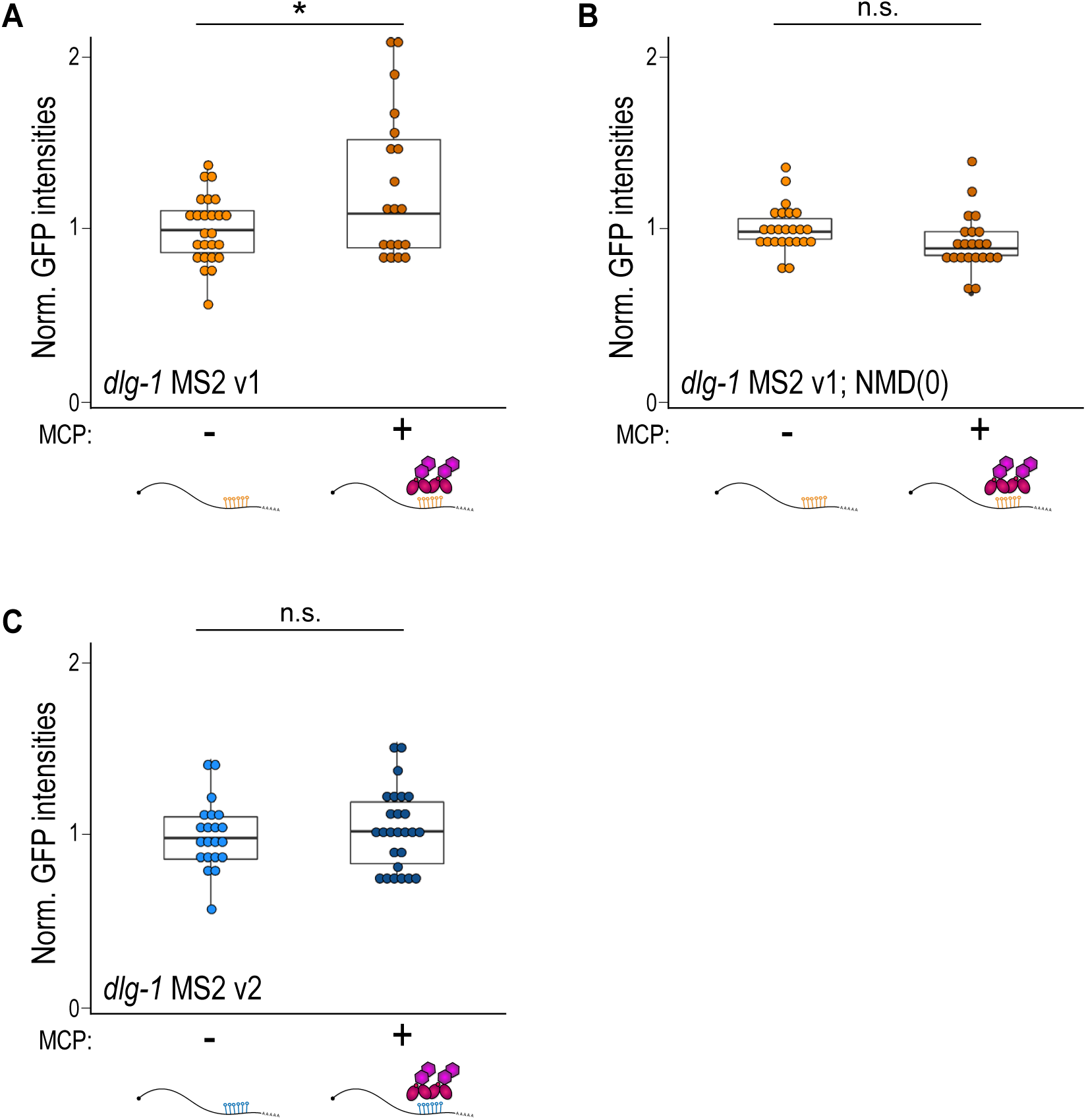
The presence of MCP marginally affects protein output in *dlg-1* MS2 lines. **A-C.** Dot plot with box plot: each dot represents the normalized GFP intensity of pharynges (*dlg-1*-tagged strain) of young adult animals (24 hours after the L4 stage) with MS2 v1 (orange) or v2 (blue) 3’UTRs in wild-type (**A**,**C**) or NMD-deficient (**B**) background in the absence (“-”) or presence (“+”) of MCP. Raw data from the minus are the same as in Fig. 1F (panels (**A**) and (**C**)) and Fig. 2F (panel (**B**)) as derived from the same experiments. Significance of statistical analyses (*t*-test, two tails): n.s. > 0.05; * < 0.01.

